# Removing bias in sequence models of protein fitness

**DOI:** 10.1101/2023.09.28.560044

**Authors:** Ada Shaw, Hansen Spinner, June Shin, Sarah Gurev, Nathan Rollins, Debora Marks

## Abstract

Unsupervised sequence models for protein fitness have emerged as powerful tools for protein design in order to engineer therapeutics and industrial enzymes, yet they are strongly biased towards potential designs that are close to their training data. This hinders their ability to generate functional sequences that are far away from natural sequences, as is often desired to design new functions. To address this problem, we introduce a de-biasing approach that enables the comparison of protein sequences across mutational depths to overcome the extant sequence similarity bias in natural sequence models. We demonstrate our method’s effectiveness at improving the relative natural sequence model predictions of experimentally measured variant functions across mutational depths. Using case studies proteins with very low functional percentages further away from the wild type, we demonstrate that our method improves the recovery of top-performing variants in these sparsely functional regimes. Our method is generally applicable to any unsupervised fitness prediction model, and for any function for any protein, and can thus easily be incorporated into any computational protein design pipeline. These studies have the potential to develop more efficient and cost-effective computational methods for designing diverse functional proteins and to inform underlying experimental library design to best take advantage of machine learning capabilities.

## Introduction

Computational protein design is an important tool for generating fit, diverse, and multi-functional proteins for therapeutics and industrial applications, as it can be intractable to experimental screen for all desirable traits across a large set of sequences. Developing machine learning methods that can minimize experiments and generate designs with desired features is key to this approach. For many real-world use cases, such as designing enzymes with novel functions or AAV capsids diverse enough to evade the immune system ^1–5^, it is essential to generate designs with high mutational depth. Yet, in contrast to this desire, predictive evaluation of protein design models is biased for deep mutational scans (DMSs) with only single mutants or else compares prediction only within single mutational depths^3,4,6–9^. While there is a challenge in evaluating such methods, which have hindered their development, it is imperative that such models be able to accurately identify high-performing designs at varying mutational depths.

Currently, unsupervised methods trained on natural sequences have been surprisingly successful at learning from evolutionary sequences to predict variant effects. ^5,6,8–15^. Yet, these are often tasks involving single mutations or at single mutational depths. However, average unsupervised model scores drop as you increase mutational depth, which while the function of a protein also drops significantly as mutations accumulate with respect to wild-type ^3,16–19^, these drops may not be the same. As an increasing percentage of synthetic sequences further away from wild-type reference sequences are non-functional^2,17–25^, good variants would have difficulty being discovered since the model severely biases for scores that are closer to the known reference sequences. Yet, since the search space of combinatorial mutants is so large, the potential positive sequences yet to be discovered in this distant search space are plentiful^16,26^, so severely biasing for spaces closer to the reference sequences closes the door on potentially very fit distant sequences.

In this paper, we demonstrate that a mutational score shift is a common model pathology across a diverse sampling of proteins that hinders precise comparisons across mutational depths. To improve the model evaluation of sequences across mutational depths, we introduce a de-biasing of scores that normalizes model scores to their background distribution of scores at each mutational depth. This method can improve our ability to pick out high-ranking sequences across mutational depths vis a vis their percentile within their background distribution. This method is generally-applicable to any unsupervised fitness prediction model, and for any function for any protein, and can thus easily be incorporated into any computational protein design pipeline and can inform experimental design.

Moreover, we demonstrate re-calibration improves semi-supervised models which combine models on evolutionary sequences with limited experimental data in novel computational tasks. Namely, we showcase how to use deep mutational scanning data around one ortholog to predict another. This task is motivated by the desire to use minimal data to extrapolate and minimize iteration between computational and experimental work, and they require properly calibrated models for mutational depths. As a proof of concept, we demonstrate a recalibrated semi-supervised model that can competitively predict the phenotype of distant orthologs in green flourescent protein (GFP) and indole-3-glycerol-phosphate synthase (IGPS) proteins.

## Results

### Experimental results show that state-of-the-art unsupervised models significantly underscore higher mutational sequences beyond their expected performance

All the state-of-the-art unsupervised protein models express the mutational score shift across most proteins. We conducted experiments on 76 proteins spanning different domains, cellular locations, and enzymatic functions (Fig S1) by generating random mutations at varying edit distances from wildtype (Figure 2A) and scoring their mutants with unsupervised models^6,10–12,27^. We observed a decrease in protein scores with increasing edit distance. However, the severity of the shift varied from protein to protein (Fig S3. In local alignment models, the severity of the shift from protein to protein was linked to less site-wise conservation. We found that lower column entropy alignments were correlated to a greater mean shift in scores for local alignment models when comparing sequences 1 mutation away to sequences 5 mutations away from wildtype (*r*2 = 0.75 EVH and *r*2 = 0.58 EVE, Fig 7). We observed that this relationship was not present in large language models. This could be due to multi-site epistasis, effects of alignment-free methods, or information from other regions of non-local protein space. Interestingly, we did not see a relationship between local sequence alignment conservation and score shifts in Tranception, despite its retrieval on the local landscape.

**Figure 1.**
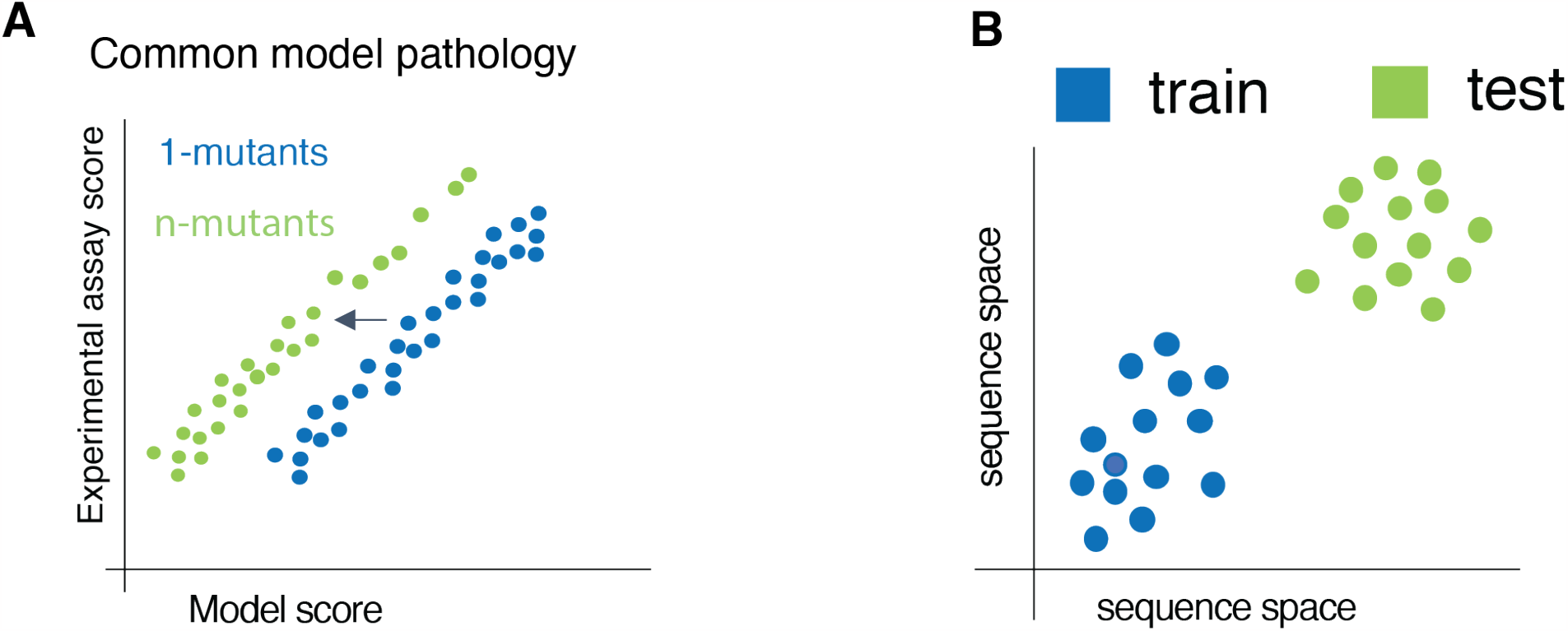
Our method addresses issues in iterative protein design that affect the ability to use both natural sequences and experimental sequences to design and predict protein sequences. **A)** Evolutionary model scores are biased for sequences close to wildtypes **B)** We address how to use labels outside one DMS’s scope by looking at its generalizability to orthologs

**Figure 2.**
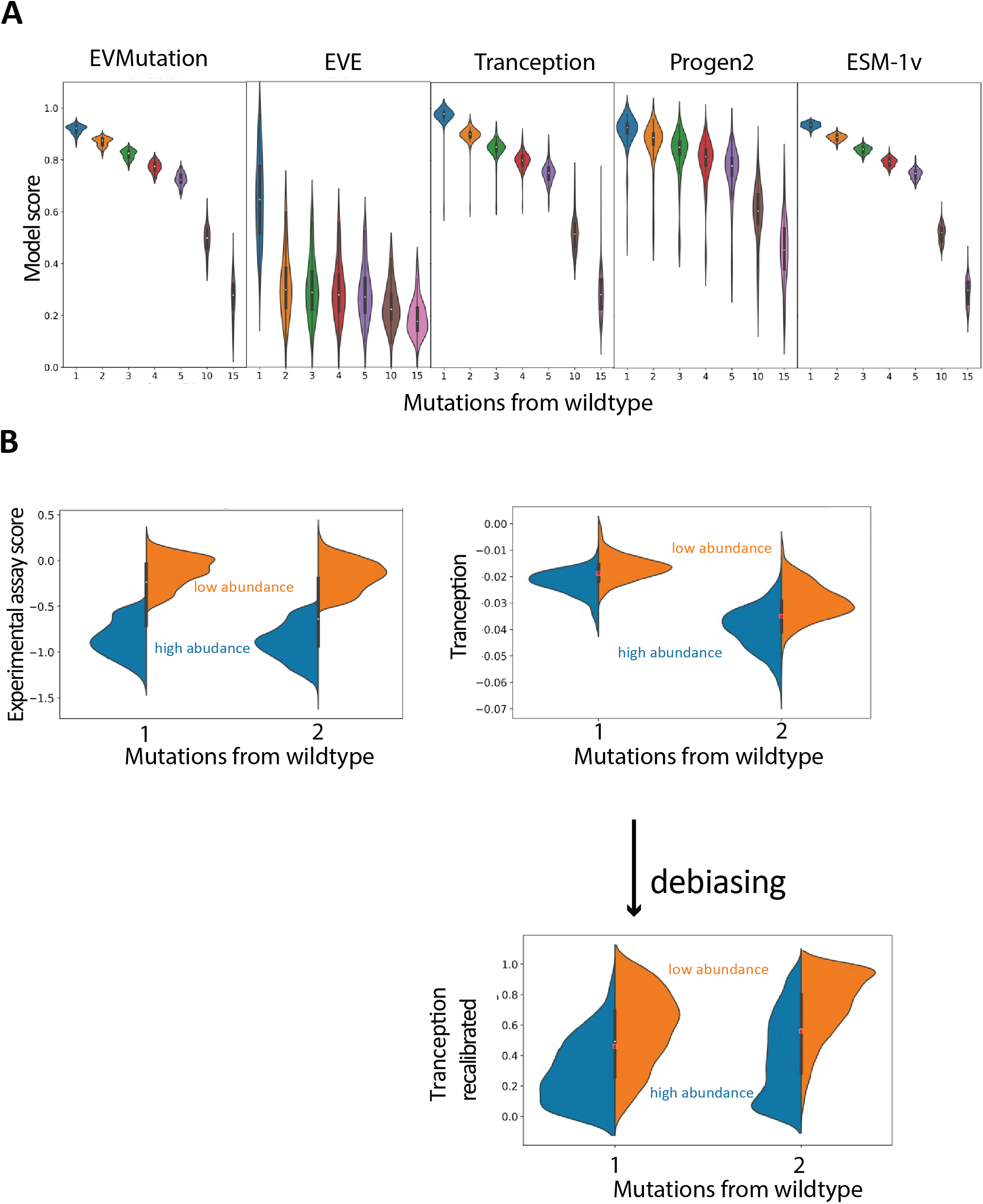
**A)** 10^4^ Randomly generated sequences for each protein were scored by the unsupervised models: EVMutation, EVE, Tranception, Progen2, and ESM1v at mutational distances of 1, 2, 3, 4, 5, 10, 15 from wildtype. Scores ranges are normalized per protein. **B)** DMS scores of protein abundance of DLG4 do not display a score shift with mutational distance from wildtype. But the model scores do (Tranception pictured here). Our recalibration method alleviates this by normalizing each mutational depth to 10^3^ randomly generated sequences at that mutational depth. We see the distributions of high and low abundance in the recalibrated scores become at a comparable range

We further investigated how this phenomenon of score shifts affects the accuracy of mutant effect prediction when experimental data characterizing multiple mutation variants are available, as shown in Table S1, which covers enzymatic proteins. We observe that the sequence scores overlap between depths and bimodal relationships in protein phenotype measurements. This is a common trend observed across all of the proteins we studied (Fig 2). This has been noted as common in protein-function relationships, in general,^17,28^. As a result, a score that indicates functional protein activity at a depth of 1 mutation may not correspond to the same level of function at a depth of 2 mutations, as seen in DLG4 where the shift is almost non-overlapping between the EVE scores (Fig 2B). This pathology is what our recalibration method aims to address. Therefore, providing specific information on how these bimodal thresholds are defined and their impact on the interpretation of scores across different mutational depths would help clarify the effect of our recalibration method in improving score accuracy.

### Percentile recalibration to background mutations improves model thresholding

The percentile recalibration is a method that calculates the scores as a percentile within the background distribution of scores at that mutational depth. It should be noted that it does not need experimental data to accomplish. With the normalization of model scores to the background distribution of scores, we observe an overlap of scores rather than a drastic mutational shift (Figure S2). To ensure that this method is not merely inflating the protein scores without improving the thresholding, we evaluated the fitness classification of 14 proteins using different models while evenly stratifying our evaluation datasets for fitness and distance from wildtype. We found that percentile recalibration generally improves the AUC across the 11 proteins (Fig 3). Specifically, the EVMutation, EVE, ESM1v, and Tranception models show a statistically significant (p-value < 0.05) improvement in protein classification with a mean AUC improvement of 4.6, 2.93, 2.65, and 2.74, respectively. While the progen2 model does not exhibit significant improvemen (p = 0.06), Progen2 does have an improved mean sample score with recalibration (mean improvement 2.063 AUC). Furthermore, at the per-mutational level, Progen2 was also not the top-performing model (being the best model for only 2 out of 14 proteins) (Fig S5). The top-performing unsupervised model across both per-mutational evaluation and sampled evaluation was found to be EVMutation which performed better on 7 out of 14 proteins. It consistently outperformed even large language models trained on the entire protein universe. However, it is worth noting that this result may be partly influenced by the use of GFP datasets, which strongly favor EVMutation. When the GFP datasets were excluded from the analysis, EVMutation remained the top-performing model for 3 out of the remaining 7 protein datasets (Fig 3A, Fig S5).

**Figure 3.**
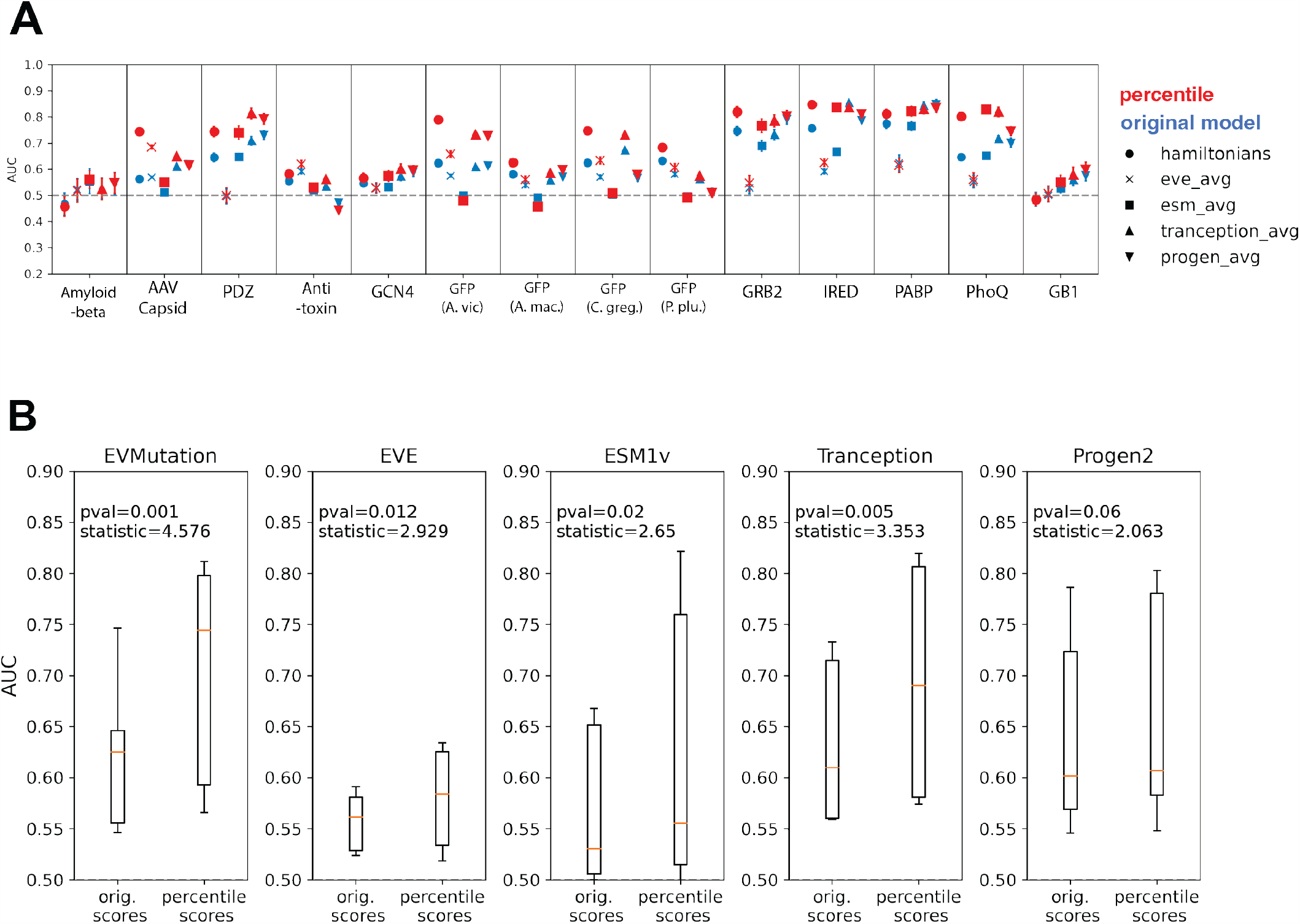
A) The 14 proteins and the improvement of the recalibration per protein. Red denotes the recalibrated AUC and blue denotes the original score. B) the improvement per model for recalibration versus the original model score for all the proteins combined.

Although EVMutation is a highly competitive model in comparison to newer sequence language models such as ESM-1v, Progen2, and Tranception, it has the highest mutational depth bias and thus has reaps largest gains from this mutational depth score de-biasing.

We note that while this evaluation does explain improved thresholding, it does not account for the functional decline with respect to distance from wild-type. To achieve that we investigate how well the method can improve the recovery of functional mutants in a functionally sparse regime and also when the top mutants are more than 2 mutations away from wildtype.

### Lower bound on functional decline using single mutants improves recovery of functional distant sequences

Here we have an example of an exhaustive deep mutational scan up to four mutations away from the wild-type of the regulator of an E. coli PhoM regulator/ PhoQ sensor pair at four different sites of the protein-protein binding interface. The functional decline of the mutants with respect to distance from *E. coli* PhoQ wild-type is a precipitous drop – with 5% of the 100 2-mutants being functional and <1% of the 150757 sequences 3 and 4 mutations away being functional. However, these 1542 functional sequences constitute 93% of the functional mutants. This makes the exhaustively scanned PhoQ system an ideal candidate for testing the efficacy of our method at identifying functional variants far away from wild-type.

Given the binary binding deep mutational scan assay on the 76 possible single mutants, we can estimate the percentage functional mutants using the non-epistatic power law (Equation 1). By correcting the sequences using percentile and then selecting top sequences using the estimated percent functional, we greatly improve the recovery of distant functional sequences at greater than 2 mutations away. Using the same threshold set at 1 mutation away and accounting for the estimated functional decline, we recover 23% of the 1542 functional sequences from the 157757 total sequences at those depths as opposed to only 3% with the uncalibrated original model scores. Here, all the models exhibit improvement with the functional decline recalibration, but only EVE does not exhibit improvement with the percentile recalibration (Fig 4).

**Figure 4.**
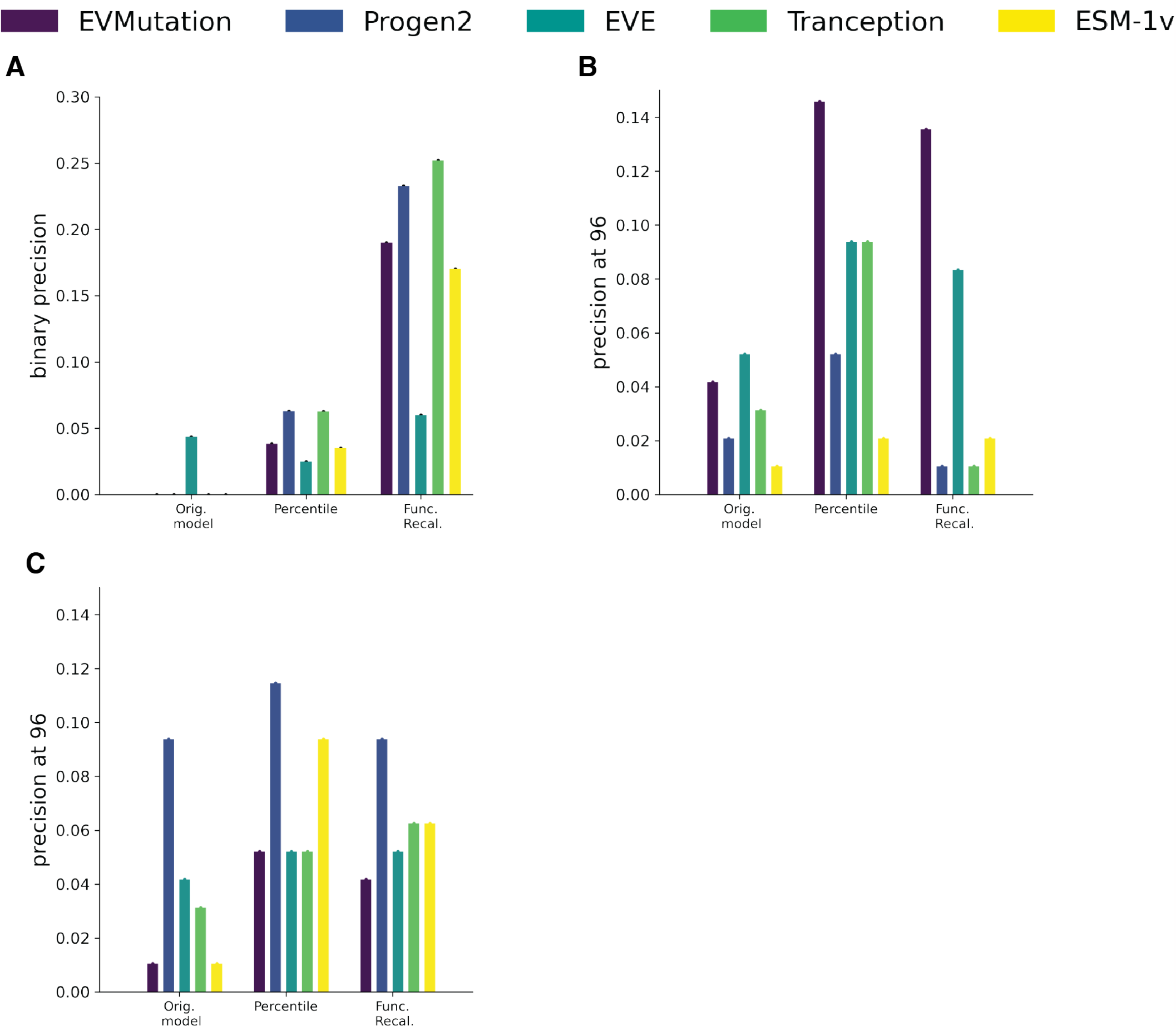
Comparison of performances for original model scores, percentile scoring, and functional de-biasing. Performed on EVMutation, Progen2, EVE, Tranception and ESM-1v **(A)** Precision improvement at greater than 2 mutations from PhoQ wild-type **(B)** Precision @ 96 sequences for anti-toxin **(C)** Precision @ 96 sequences for GCN4

Although the PhoQ dataset is very comprehensively sampled. Its experimental measurement is strictly binary therefore it does not allow a close look at how well these recalibrated models can pick out high-performing mutants.

### Precision @k metric measures model precision at collecting top-performing proteins

Here, we focus on examining biases related to sequence rankings and compare them to an experimental setup that employs a continuous score. When evaluating sequence rankings, our primary metric of choice is precision @ k. We favor this metric because it prioritizes the retrieval of top-performing sequences, distinguishing it from the commonly used Spearman’s *ρ* metric. While precision @ k doesn’t explicitly give preference to sequences from extensively sampled depths, it inherently leans towards identifying superior mutants within depth ranges that have more samples (Fig S8).

An ideal dataset to validate this method would have a more egalitarian sampling of sequences at each depth but as a dataset does not exist we focus our evaluation where the sampling of sequences at higher depths is more comprehensive than that at lower depths. Unlike most datasets typically which offer only limited samplings of potential combinatorial variants in comparison to lower mutational depths, PhoQ, anti-toxin, and GCN4 datasets, are more comprehensive at higher mutational depths. We note that although they are more comprehensive it does not necessarily mean that it covers a greater percentage of the possible mutants at that mutational depth.

We investigate GCN4 and the anti-toxin deep mutational scans primarily because the datasets are not both biased for lower mutational sequences and gives us an opportunity to see if the model can pick up distant highly ranked sequences that would otherwise be marked non-functional. Yet, the significant bias in sampling towards sequences with 4-5 mutational differences raises a concern it may also yield skewed results due to this sampling bias. We choose precision @ 96 to simulate the experiments one can run on a 96-well plate. For all these datasets, 96 sequences represent <5% of all the total sequences of each of the datasets (Table S1). Here we observe that recalibrating to a background distribution improves the recovery of all of these sequences for both datasets but little gain from using the functional decline. For the anti-toxin, EVMutation leads the recovery with 14.6% of the top 96 sequences recovered using the percentile method (versus at best 5% using any model’s original score). For GCN4, Progen2 modestly improves the recovery with 11.4% sequences recovered using percentile and 9.4% sequences recovered using the original score.

### A semi-supervised model tracks top SOTA supervised or semi-supervised models on orthologs

(Hsu et al. 2022) show that their method of combining evolutionary models with labeled data improves the predictive capability when training on small subsets of functional assays and extrapolating to predict held-out sequences with mutational depths similar to those in the training set^7^. We show that this model sometimes struggles to outperform linear regression when trained on a DMS to predict distant orthologs. This implies that although the evolutionary information allows the model to impute within the distances of the training set, it has difficulty using that information to extrapolate to more distant variation on the background of an ortholog. This is pertinent when designing sequences in orthologous proteins given a DMS. We ameliorate this issue using an alternate scheme to combine models’ evolutionary data and experimental data – the product of experts, where the probability distribution is a product of simpler distributions^29^. In our case, our two distributions are the logistic regression model multiplied by our recalibrated evolutionary models. We refer to this model as the PoE method. To evaluate the performance of this semi-supervised model, we compared it to other supervised methods and the augmented Potts. We examined two datasets: the multi-mutational deep mutational scans around each of the four green fluorescent protein orthologs^17^ and the single-mutational scans around each of the three 3-indole phosphate glycerol synthase (IGPS)^30^.

For the green fluorescent protein (GFP) dataset, we trained on the 51k sequences in the multi-mutational deep mutational scan of Aequorea victoria (avGFP) and predicted other multi-mutational deep mutational scans of the other orthologs of GFP, including Aequorea macrodactyla (amacGFP), Clytia gregaria (cgreGFP), and Pontellina plumata (ppluGFP). amacGFP, cgreGFP, and ppluGFP are at from 82, 41, 18 percent similarity from avGFP. Performance was measured using the area under the receiver operating characteristic curve (AUC) when evenly sampling all the mutational distances and binary cases using the sampling technique described in the methods.

The PoE method that uses the recalibrated Potts score performed at the same level or moderately outscored the augmented Potts and other supervised methods when scoring distant orthologs. It was trained on one ortholog to predict another. Specifically, when predicting amacGFP sequences at 82% similarity, both semi-supervised methods performed on par with a simple linear model at 0.75 AUC. When predicting cgreGFP at 41% similarity, our semi-supervised model barely outperformed all the models at 0.8 AUC (vs. 0.79 AUC of the linear model runner-up). The higher performance of the avGFP-trained models on cgreGFP was also observed for the unsupervised Potts where training on sequences centered about avGFP performed better on cgreGFP versus amacGFP (0.68 AUC versus 0.58, respectively). When predicting ppluGFP at 18 percent similarity, the Potts model outperformed all the models. This suggests that evolutionary data is more important than the training labeled data at that mutational distance from the labels (Fig 5A).

**Figure 5.**
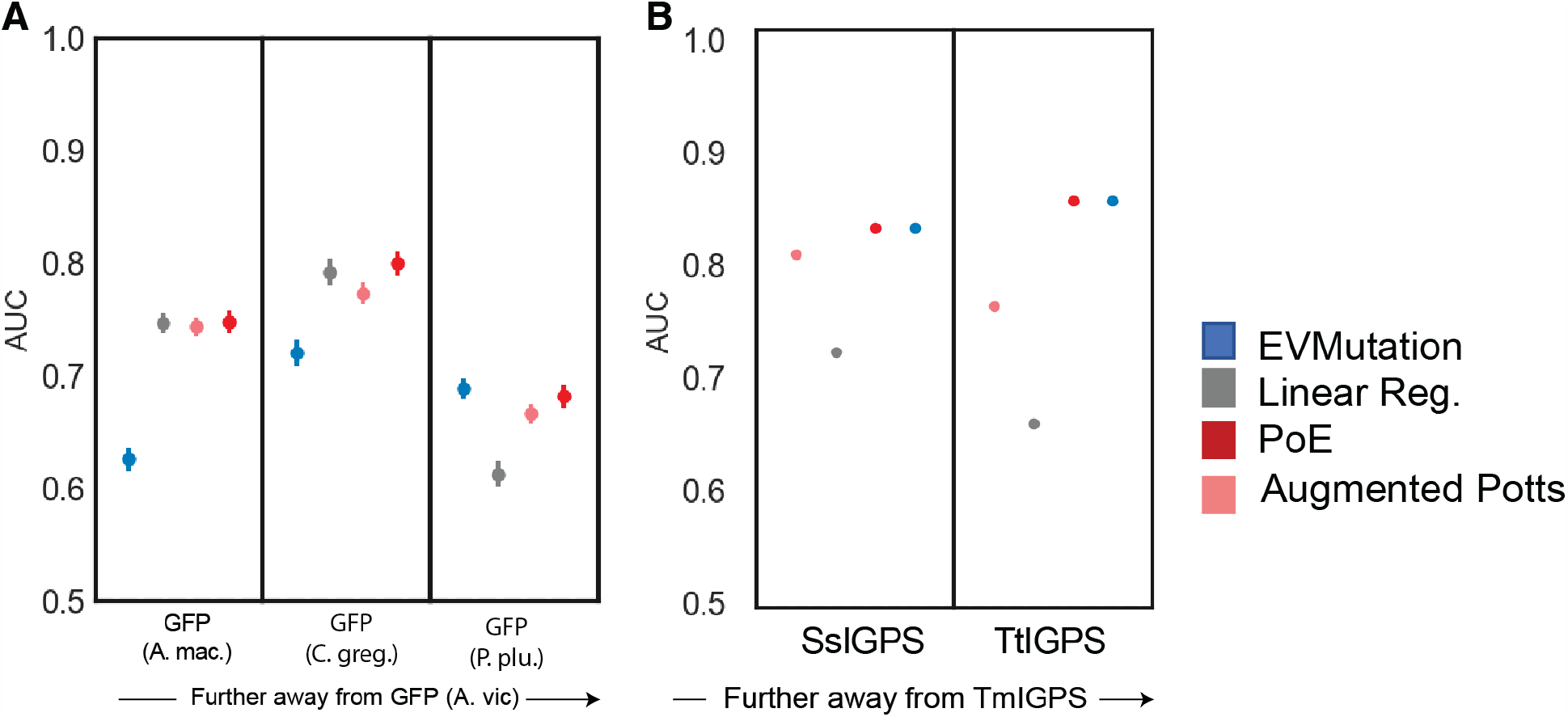
Model performance measured in AUC for 4 top performing models on predicting orthologs given one DMS. **(A)** When training on Aequorea victoria (avGFP) and predicting Aqueous macrodactyla (amacGFP), Clytia gregaria (cgreGFP), and Pontellina plumata (ppluGFP). **(B)** When training on T. maritima (TmIGPS) and testing on S. solfataricus (SsIGPS) and T. thermophilus (TtIGPS).

For the IGPS dataset (Fig 5B), we trained on one IGPS (1600 single mutants) to predict the other ortholog proteins (1600 single mutants per ortholog). The IGPS proteins were approximately 30% similarity to each other. We evaluated the model performance using the AUC of the model with respect to a binary label. In all cases, our semi-supervised model and Potts model outperformed supervised methods and augmented supervised models. The semi-supervised model closely tracked the unsupervised Potts model without exceeding its performance.

Lastly, we observed no advantage of large language models trained on the global protein-verse as well as fine-tuned on the local alignment compared to models using local alignment alone for both the GFP and IGPS datasets.

## Discussion

Current methods that leverage evolutionary protein sequences to predict variant fitness are known to downweight sequences further away from wildtype. The model will automatically consider sequences further from wildtype as worse than sequences closer to wildtype. This complicates the chances of identifying successful protein candidates, especially if sequence diversity is a criterion for success. The percentile recalibration method we introduce improves recovery of top-performing protein sequences for multi-mutant datasets. It also prevents possibly fit proteins in further mutational depths from being classified as non-fit.

While recalibrating sequence scores given a sparse natural sampling is of potentially great use for protein design efforts, existing datasets that span large ranges in mutational depths are rare because of challenges in effectively expressing and screening a range of natural variants in appropriate reaction conditions. The data available for validation of our approach is limited as there are many more single variant datasets than multiple variant datasets. Though altogether we analyze 14 proteins that represent 7 distinct phenotypes, this dataset is heavily biased toward a single protein, GFP. The ortholog prediction analysis is similarly limited to only TIM barrel proteins: IGPS and GFP. As the importance of DMSs across mutational depths becomes clearer, and more data becomes available, our de-biasing approach has the potential to have a greater impact. As a result of the current data limitation, we do not observe significant benefits from the functional decline de-biasing strategy from many of the existing datasets. Because of the present sampling bias for low mutational depths and experimental noise, it is difficult to assess whether our functional decline de-biasing method is worse at picking at top mutants or whether the datasets are limited. More fine-tuned recalibration methods would likely require experimental information since these differences can often be more specific to the experiment and not the protein’s natural fitness itself. Methods that can incorporate epistasis and increases in standard deviation into the functional decline can also improve the functional decline estimation.

Nonetheless, this large-scale analysis provides important insights into unsupervised modeling and experimental approaches to best learn about sequences at high mutational depths. We provide a method to better compare model scores across mutational depths – making it easier to identify high-performing mutants in higher mutational depths. We show how this is especially pertinent in situations of very sparse functional sequences. We demonstrate how this recalibrated model can improve the generalizability of the model, allowing it to be incorporated into semi-supervised methods that improve the search for functional sequences in distant orthologs.

We identify a common pathology in sequence models trained on evolutionary sequences and we provide an ad-hoc method that can be applied to any model can improve the search for functional proteins – whether it be enzymes, antibodies, or capsid proteins. This method can, without additional experimental data, improve the ability to identify functional sequences at distances from wildtype. Moreover, we present a lightweight semi-supervised method specifically designed for identifying functional distant orthologs of GFP and IGPS proteins, which outperforms the current state-of-the-art semi-supervised approach.

## Methods

In this study, we evaluated the performance of several unsupervised machine learning state-of-the-art models on diverse publicly available datasets (Table S1)^10–12^. We developed a novel method for improving the accuracy of these models by recalibrating their scores with respect to the background distribution of random mutations at different mutational depths. Additionally, we evaluated the generalization ability of supervised and semi-supervised models using several datasets to test model generalization across orthologs. In this section, we present the methods we used to train, recalibrate, and evaluate our models.

### Model generation

#### Multiple sequence alignments

Multiple sequence alignments of evolutionary sequences were constructed using jackhmmer^31,32^, an iterative profile-HMM-based search tool, against the uniref100 database. We optimized the search depth to maximize sequence coverage and the effective number of sequences included after clustering similar sequences as previously reported^10–12^ and to optimize contact map accuracy. PDB structures that were used to assess the contact map are denoted in Table S2.

#### Models

We evaluated Progen2, Tranception, ESM1v, EVMutation, and EVE as a representative sample of state-of-the-art unsupervised models encompassing masked language models, autoregressive language models, maximum entropy models, and variational autoencoder methods.

### Bias removal

#### Percentile

To normalize the changing standard deviation per mutational depth, we generated background mutations for each protein in Table S1 at all mutational depths and then calculated the percentile of each score relative to the corresponding background distribution.

#### Estimate of functional decline

To show the functional decline in mutants further away from wild type, the top percent functional mutants per mutational depth *P*_*additive*_(*n* = *N*) were approximated using a power law that had a no epistasis assumption:

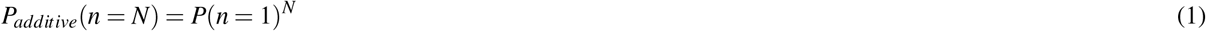

where *P*(*n* = 1) is the fraction of single mutants that are fit and *N* is the mutational depth where we are approximating the percent functional. Figure 9 shows all the empirical data percent functional versus the *P*_*additive*_. It should be noted however, datasets with the large deviations from the *P*_*additive*_ trend, i.e. GCN4, IRED, AAV (designed) are non-randomized mutagenesis. Additionally, these empirical percentages can be extremely variable in situations where there is a low number of sequences at that depth. *Continuous score method:* The continuous score is a *de facto* probability of being in the functional group. First, we calculate *μ*_*n,null*_, *σ*_*n,null*_, the mean and standard deviation of the null distribution’s top *P*(*n* = *N*) percentiles for each depth, *n*. Then, we fit normal cumulative distributive function to *μ*_*n,null*_, *σ*_*n,null*_. The top *P*(*n* = *N*) percentile sequences in the from the deep mutational scan are then standard scaled using *μ*_*n,null*_, *σ*_*n,null*_ and the cumulative distribution function is calculated for those scores.

#### Binary method

We mark the top *P*_*additive*_(*n* = *N*) percentile of sequences as functional.

### Evaluation

#### Experimental data

We acquired publicly available multi-mutant data 1. For binarization into functional or non-functional scores, if the data was visually bimodal, we fit a 2-component gaussian mixture model to the scores and if the data was exponential, we followed the protocols as defined by Starr et al., 2020^33^.

#### Data sampling to evaluate performance at different mutational depths

The scores often have a drop in functionality with respect to distance, which compounds with the model’s propensity to poorly rank distant sequences, forming a Simpson’s paradox. We conducted a sampling and evaluation process to mitigate the potential biases introduced by the selection of evaluation samples based on cases or mutational depths. Specifically, we evaluated each model’s ability to accurately classify AUC across different mutational depths by evenly sampling cases (determined via either the median or mixture model, depending on the dataset). This was done for each depth while ensuring that each depth contained at least 100 sequences and that the case percentage was at least 15% of the lesser case in order to prevent a mutational depth sample from being over-inflated with the same sequence. We repeated this resampling process 100 times to obtain the mean and standard deviation of the AUC. Furthermore, we examined the performance of the models at each mutational depth to ensure that the top-performing model before recalibration remained the top-performing model after recalibration.

#### Precision with lower mutational depths

For certain datasets, ranking is not possible – this is either because the experimental assay only has a binary label or because the correlation of the replicates is low^16,17^. For these datasets, we evaluate the performance using the precision of the model at different mutational depths using the same threshold drawn at the single mutational depth.

#### Supervised and semi-supervised model evaluation

For evaluating the supervised and semi-supervised models, we tested model generalization across orthologs by training on one ortholog and predicting the rest using the GFP (green fluorescent protein)^17^ and IGPS (Indole 3-Glycerol Phosphate Synthase)^30^ datasets. We evaluated several supervised models, including CNN, RNN, Linear regression, and LSTM. The linear regression model was chosen as it outperformed the other models. Additionally, we tested various semi-supervised models, including augmented Potts, augmented EVE, augmented ESM1v, augmented Tranception, and augmented Progen2. We report the results for augmented Potts only, as it performed on par or outperformed all the other unsupervised augmentations across the 12 datasets. Furthermore, we developed a semi-supervised model inspired by the Product of Experts format: *Y* = *F*(*x*) × *G*(*x*), where *F*(*x*) is a supervised model trained on labels and *G*(*x*) is the recalibrated unsupervised model (if we are testing multiple depths). Notably, a sigmoid was added to the last layer of the models before multiplying as it more closely matched the shape of the bimodal phenotype distribution.

## Supporting information

DMS Dataset Information

## Author contributions statement

AYS collected data, designed the method, analyzed the experimental data, and wrote the manuscript. HS helped with data analysis and recalibration method design, JE Shin collected datasets and aided project inception, SG helped with recalibration method design and revisions, NR helped with method design, DSM oversaw the project.

## Code Availability

Data and code is available on GitHub

## Supplementary

**Table 1.**
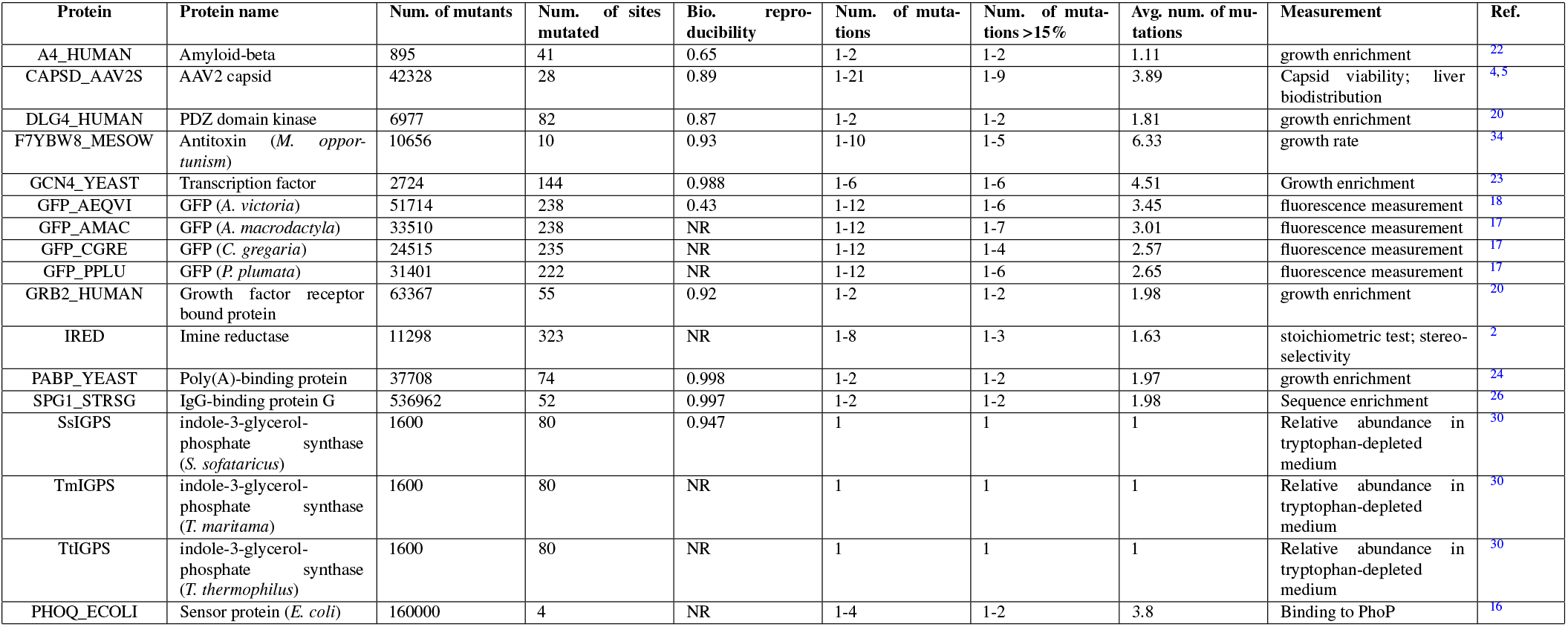
Protein deep mutational scan datasets.

**Table 2.**
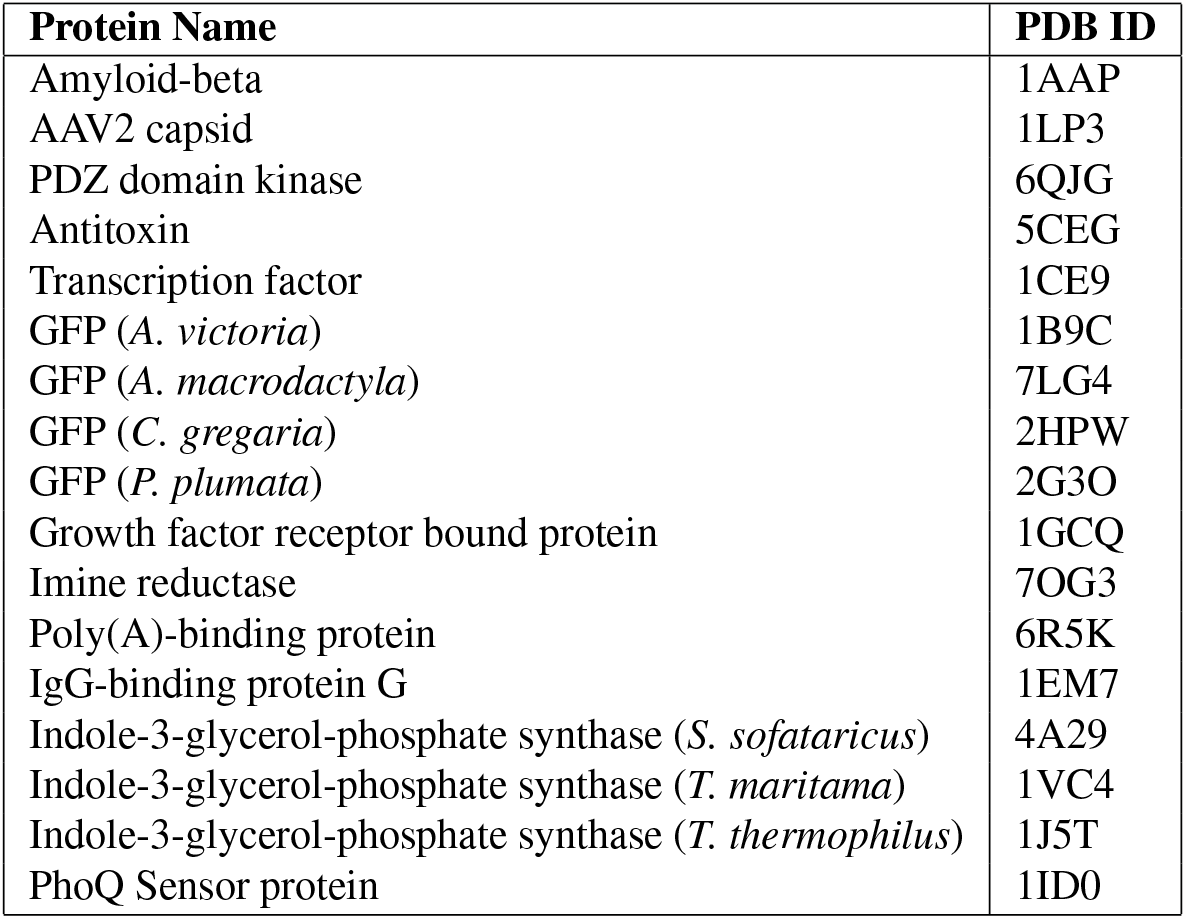
Protein Names and associated PDB IDs.

**Supplementary Figure 1.**
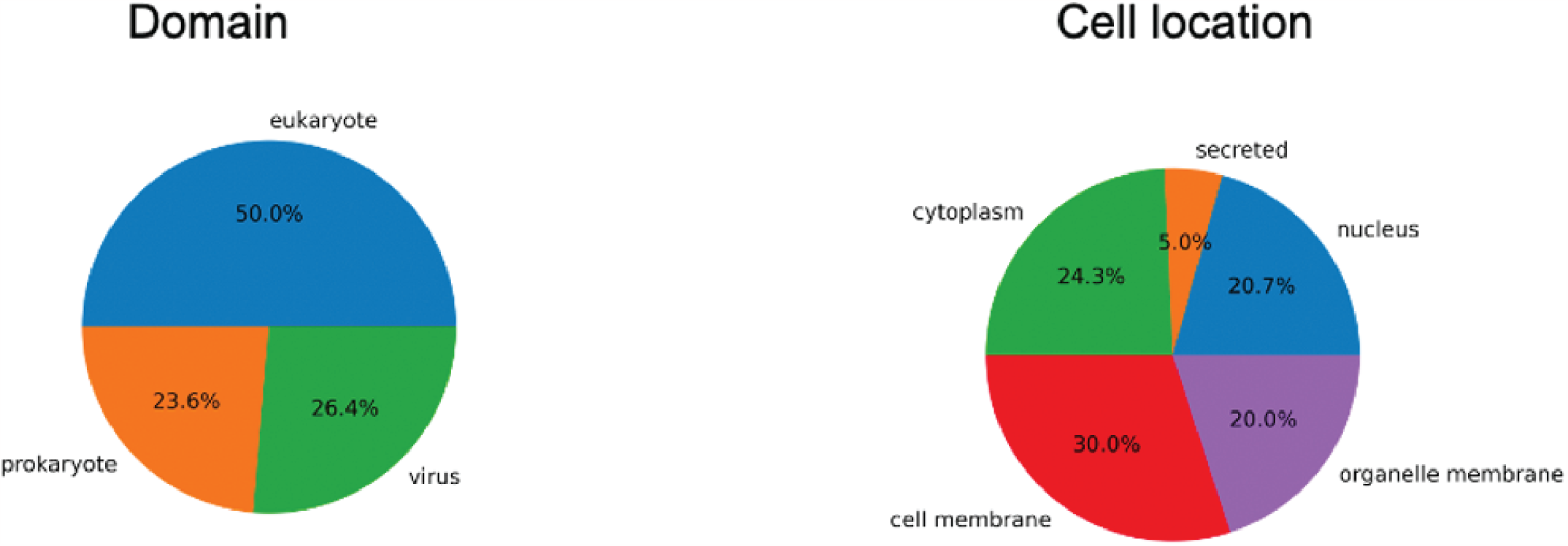
Domain and cell location of proteins used in large scale study

**Supplementary Figure 2.**
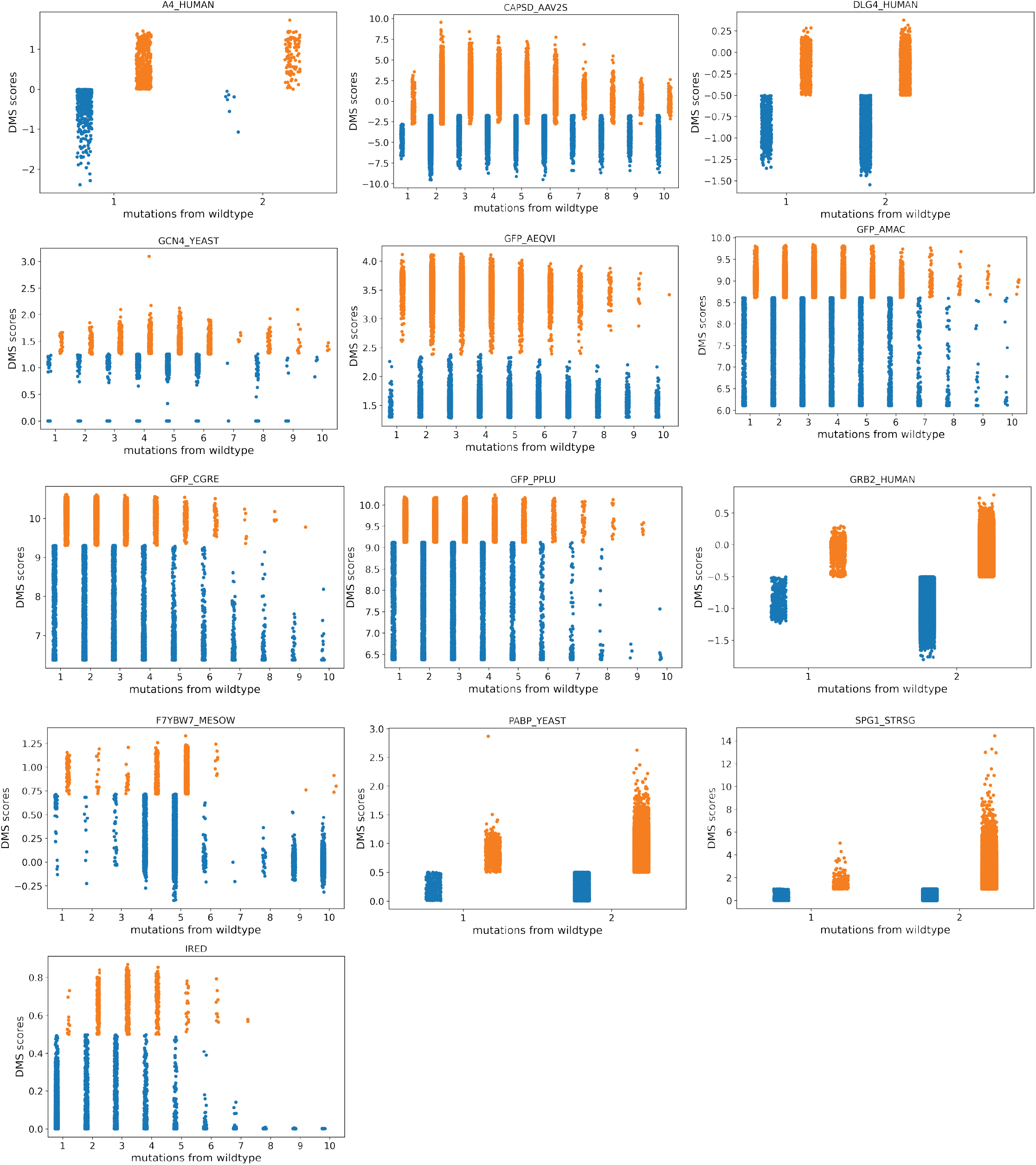
experimental assay values for each DMS. See Table S1 for full details

**Supplementary Figure 3.**
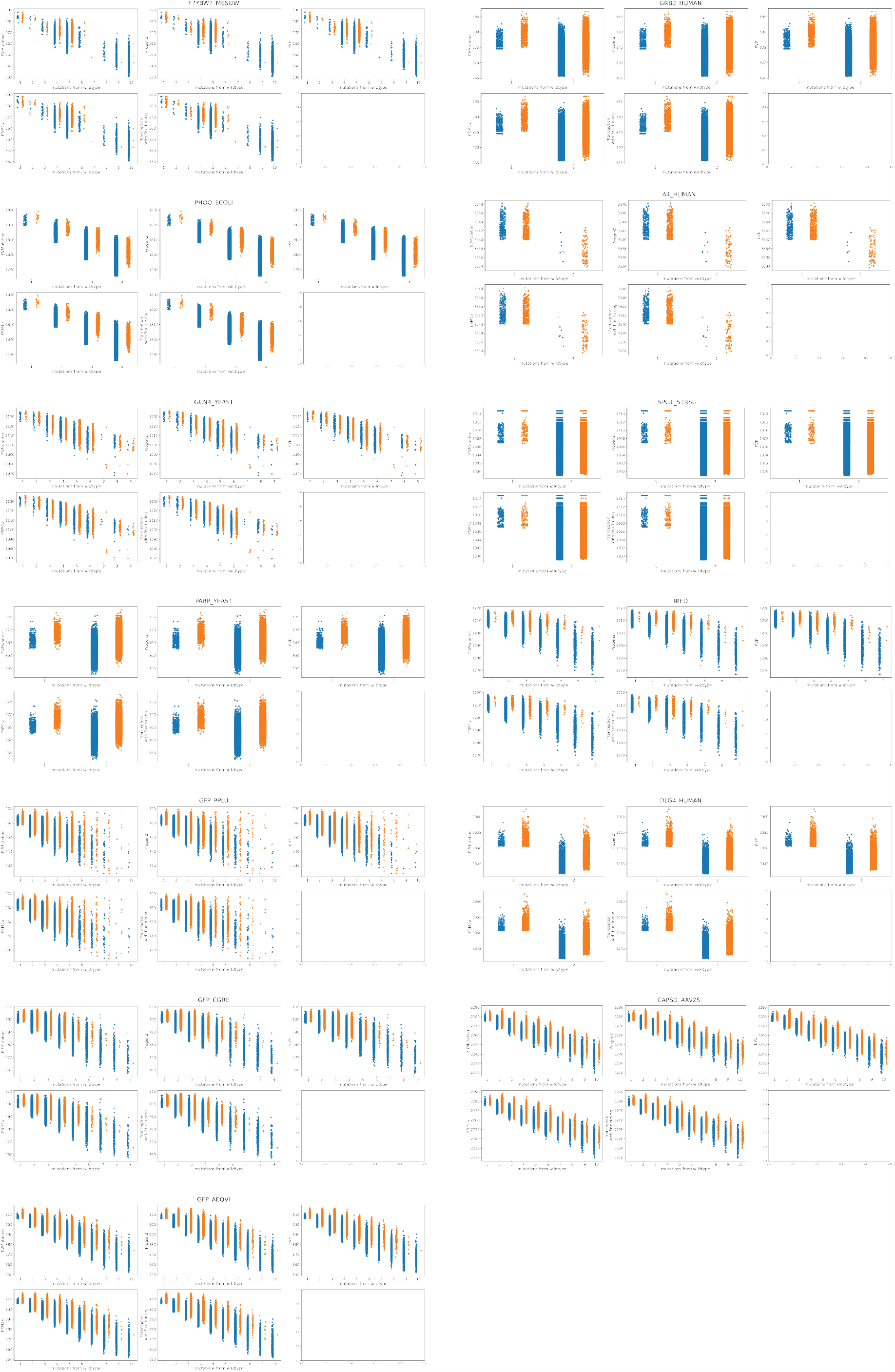
original model scores for EVMutation, Tranception, ESM-1v, Progen2, EVE. Functional sequences are denoted in orange and non-functional sequences are denoted in blue.

**Supplementary Figure 4.**
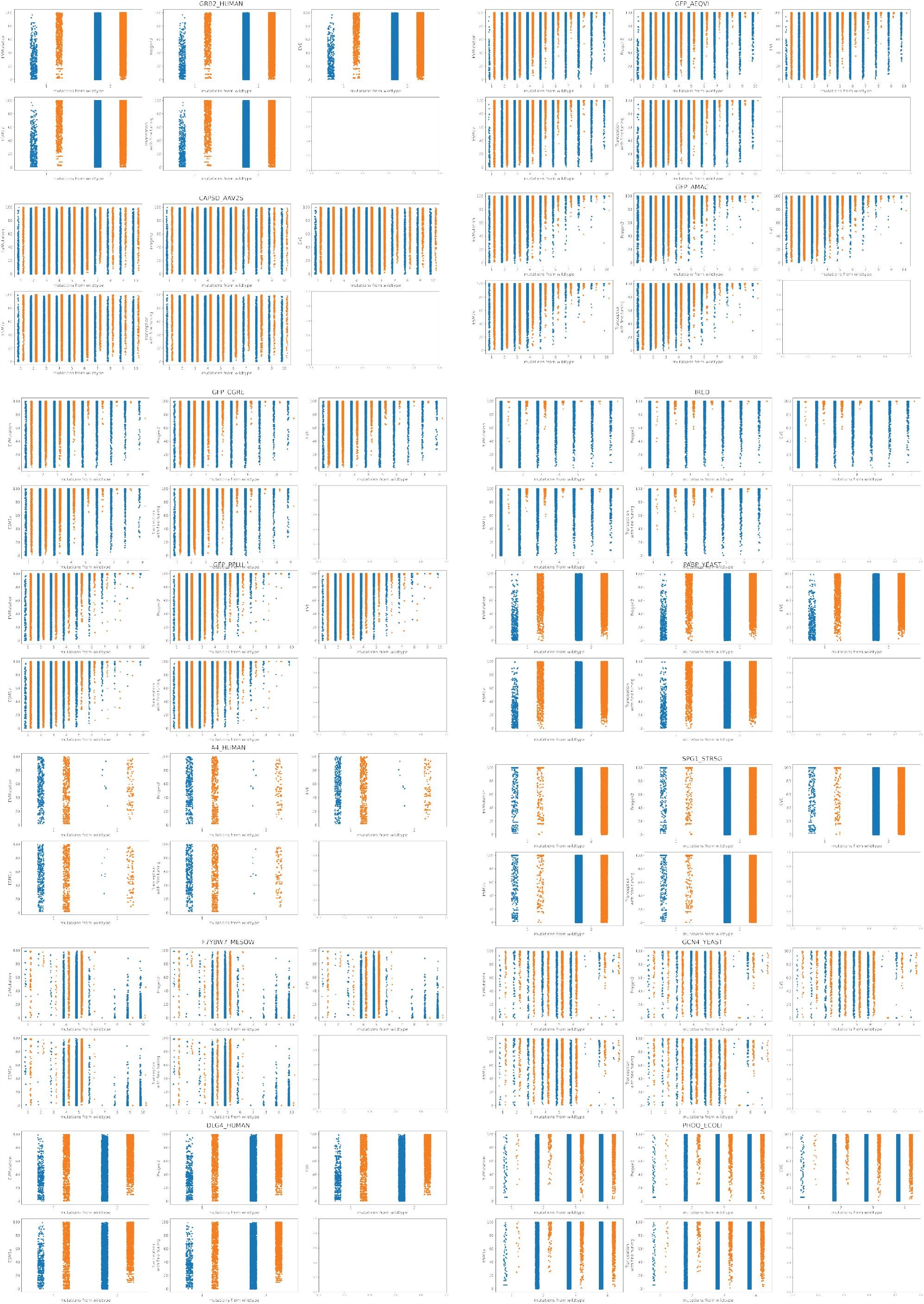
percentile model scores for EVMutation, Tranception, ESM-1v, Progen2, EVE. Functional sequences are denoted in orange and non-functional sequences are denoted in blue.

**Supplementary Figure 5.**
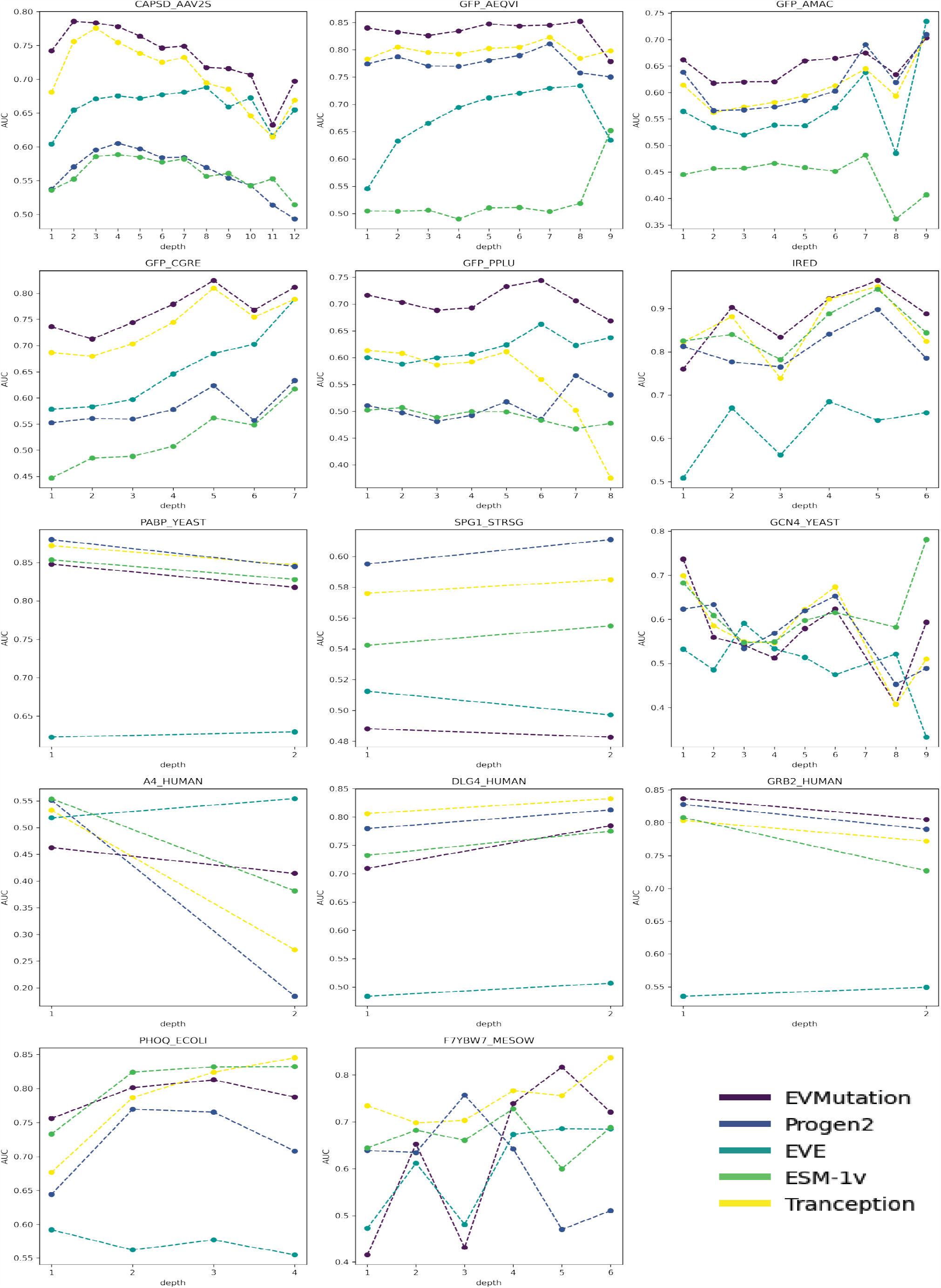
per mutational AUC for each unsupervised model

**Supplementary Figure 6.**
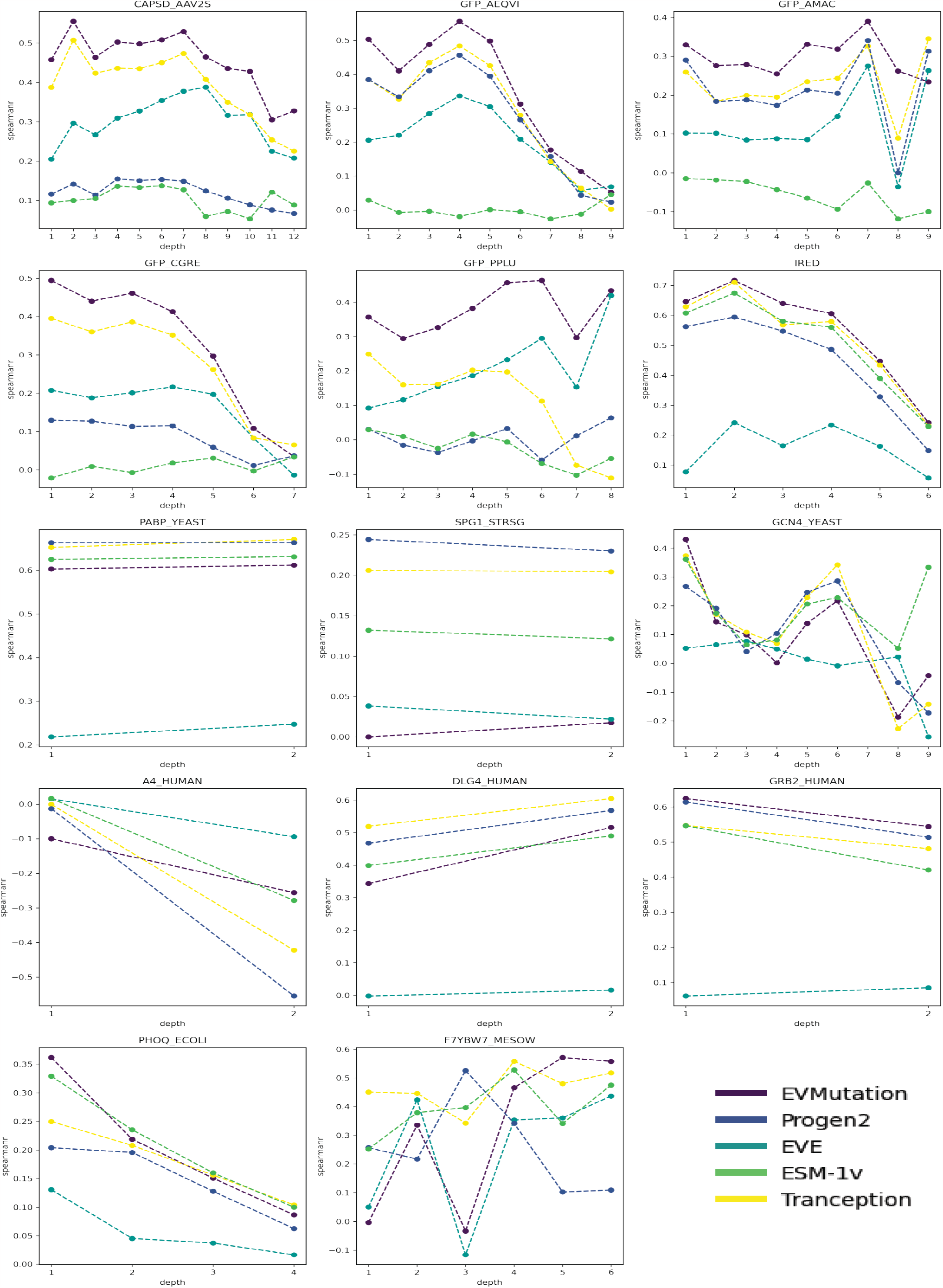
per mutational spearman *ρ* for each unsupervised model

**Supplementary Figure 7.**
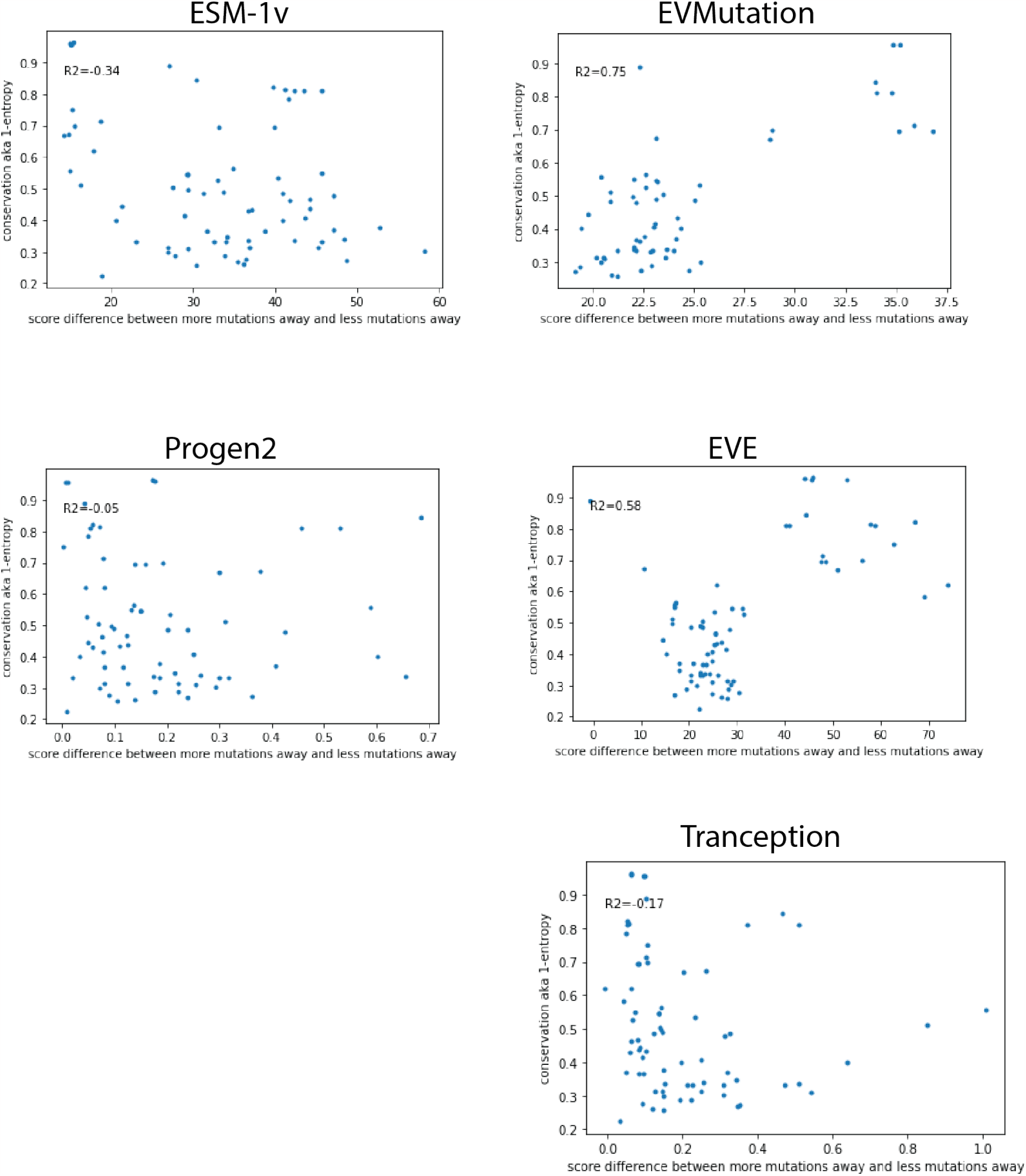
entropy vs Δ*mean*_*model*,1*muts*_ *−mean*_*model*,5*muts*_

**Supplementary Figure 8.**
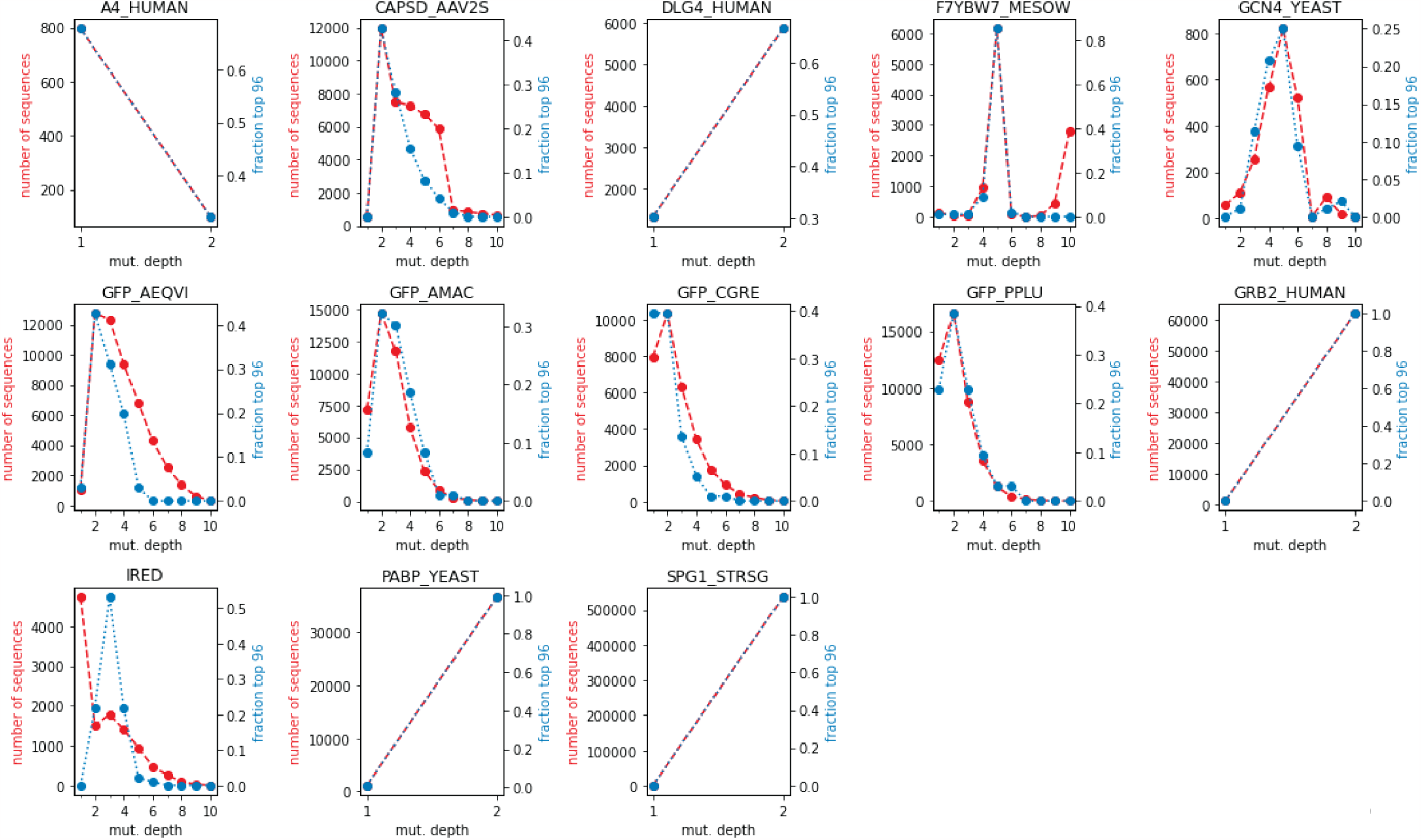
mutational depth versus fraction of top 96 (experimental performance) sequences for proteins (blue) and mutational depth versus number sequences sampled at depth (red)

**Supplementary Figure 9.**
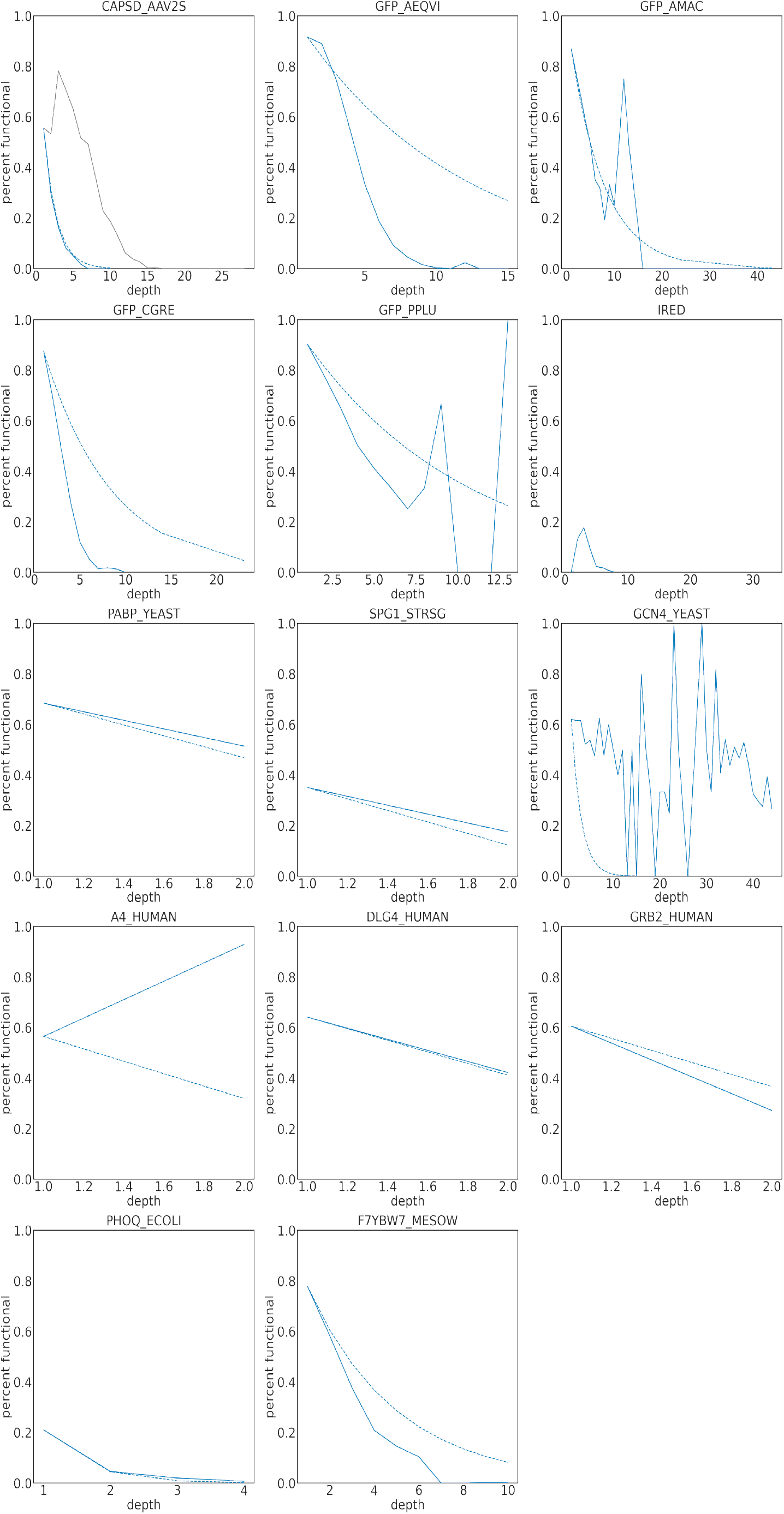
comparison of *P*_*additive*_ (dashed blue line) and *P*_*empirical*_(solid blue line) for percent functional. For AAV2S capsid, the randomized mutants and designed mutants are denoted by the solid blue line and solid gray line, respectively.

## References

1. Erickson, E. et al. Sourcing thermotolerant poly(ethylene terephthalate) hydrolase scaffolds from natural diversity. Nat. Commun. 13, 7850 (2022).

2. Ma, E. J. et al. Machine-Directed evolution of an imine reductase for activity and stereoselectivity. ACS Catal. 11, 12433–12445 (2021).

3. Ogden, P. J., Kelsic, E. D., Sinai, S. & Church, G. M. Comprehensive AAV capsid fitness landscape reveals a viral gene and enables machine-guided design. Science 366, 1139–1143 (2019).

4. Bryant, D. H. et al. Deep diversification of an AAV capsid protein by machine learning. Nat. Biotechnol. 39, 691–696 (2021).

5. Sinai, S., Jain, N., Church, G. M. & Kelsic, E. D. Generative AAV capsid diversification by latent interpolation (2021).

6. Notin, P. et al. Tranception: Protein fitness prediction with autoregressive transformers and Inference-Time retrieval. Proc. 39th Int. Conf. on Mach. Learn. 169, 16990–16999 (2022).

7. Hsu, C., Nisonoff, H., Fannjiang, C. & Listgarten, J. Combining evolutionary and assay-labelled data for protein fitness prediction (2021).

8. Alley, E. C., Khimulya, G., Biswas, S., AlQuraishi, M. & Church, G. M. Unified rational protein engineering with sequence-based deep representation learning. Nat. Methods 16, 1315–1322 (2019).

9. Madani, A. et al. ProGen: Language modeling for protein generation. (2020). 2004.03497.

10. Shin, J.-E. et al. Protein design and variant prediction using autoregressive generative models. Nat. Commun. 12, 1–11 (2021).

11. Marks, D. S., Hopf, T. A. & Sander, C. Protein structure prediction from sequence variation. Nat. Biotechnol. 30, 1072–1080 (2012).

12. Riesselman, A. J., Ingraham, J. B. & Marks, D. S. Deep generative models of genetic variation capture the effects of mutations. Nat. Methods 15, 816–822 (2018).

13. Frazer, J. et al. Disease variant prediction with deep generative models of evolutionary data. Nature 599, 91–95 (2021).

14. Marquet, C. et al. Embeddings from protein language models predict conservation and variant effects. Hum. Genet. 141, 1629–1647 (2022).

15. Thadani, N. N. et al. Learning from pre-pandemic data to forecast viral escape. bioRxiv 2022–07 (2022).

16. Podgornaia, A. I. & Laub, M. T. Protein evolution. pervasive degeneracy and epistasis in a protein-protein interface. Science 347, 673–677 (2015).

17. Gonzalez Somermeyer, L. et al. Heterogeneity of the GFP fitness landscape and data-driven protein design. Elife 11 (2022).

18. Sarkisyan, K. S. et al. Local fitness landscape of the green fluorescent protein. Nature 533, 397–401 (2016).

19. Pokusaeva, V. O. et al. An experimental assay of the interactions of amino acids from orthologous sequences shaping a complex fitness landscape. PLoS Genet. 15, e1008079 (2019).

20. Faure, A. J. et al. Mapping the energetic and allosteric landscapes of protein binding domains. Nature 604, 175–183 (2022).

21. Sinai, S., Kelsic, E., Church, G. M. & Nowak, M. A. Variational auto-encoding of protein sequences. (2017). 1712.03346.

22. Seuma, M., Faure, A. J., Badia, M., Lehner, B. & Bolognesi, B. The genetic landscape for amyloid beta fibril nucleation accurately discriminates familial alzheimer’s disease mutations. Elife 10 (2021).

23. Staller, M. V. et al. A High-Throughput mutational scan of an intrinsically disordered acidic transcriptional activation domain. Cell Syst 6, 444–455.e6 (2018).

24. Melamed, D., Young, D. L., Gamble, C. E., Miller, C. R. & Fields, S. Deep mutational scanning of an RRM domain of the saccharomyces cerevisiae poly(a)-binding protein. RNA 19, 1537–1551 (2013).

25. Ding, D. & al., E. Co-evolution of interacting proteins through non-contacting and non-specific mutations. Nat. Ecol. & Evol. 6, 590–603 (2022).

26. Olson, C. A., Wu, N. C. & Sun, R. A comprehensive biophysical description of pairwise epistasis throughout an entire protein domain. Curr. Biol. 24, 2643–2651 (2014).

27. Rives, A. et al. Biological structure and function emerge from scaling unsupervised learning to 250 million protein sequences. Proc. Natl. Acad. Sci. 118, e2016239118 (2021).

28. Kemble, H., Nghe, P. & Tenaillon, O. Recent insights into the genotype-phenotype relationship from massively parallel genetic assays. Evol. Appl. 12, 1721–1742 (2019).

29. Hinton, G. E. Training products of experts by minimizing contrastive divergence. Neural Comput. 14, 1771–1800 (2002).

30. Chan, Y. H., Venev, S. V., Zeldovich, K. B. & Matthews, C. R. Correlation of fitness landscapes from three orthologous TIM barrels originates from sequence and structure constraints. Nat. Commun. 8, 1–12 (2017).

31. Johnson, L. S., Eddy, S. R. & Portugaly, E. Hidden markov model speed heuristic and iterative HMM search procedure. BMC Bioinforma. 11, 431 (2010).

32. Eddy, S. R. Accelerated profile HMM searches. PLoS Comput. Biol. 7, e1002195 (2011).

33. Starr, T. N. et al. Deep mutational scanning of sars-cov-2 receptor binding domain reveals constraints on folding and ace2 binding. cell 182, 1295–1310 (2020).

34. Ding, D. et al. Protein design using structure-based residue preferences (2023).

